# Convergent evolution and structural adaptation to the deep ocean in the protein folding chaperonin CCTα

**DOI:** 10.1101/775585

**Authors:** Alexandra A.-T. Weber, Andrew F. Hugall, Timothy D. O’Hara

## Abstract

The deep ocean is the largest biome on Earth and yet it is among the least studied environments of our planet. Life at great depths requires several specific adaptations, however their molecular mechanisms remain understudied. We examined patterns of positive selection in 416 genes from four brittle star (Ophiuroidea) families displaying replicated events of deep-sea colonization (288 individuals from 216 species). We found consistent signatures of molecular convergence in functions related to protein biogenesis, including protein folding and translation. Five genes were recurrently positively selected, including CCTα (Chaperonin Containing TCP-1 subunit α), which is essential for protein folding. Molecular convergence was detected at the functional and gene levels but not at the amino-acid level. Pressure-adapted proteins are expected to display higher stability to counteract the effects of denaturation. We thus examined *in silico* local protein stability of CCTα across the ophiuroid tree of life (967 individuals from 725 species) in a phylogenetically-corrected context and found that deep sea-adapted proteins display higher stability within and next to the substrate-binding region, which was confirmed by *in silico* global protein stability analyses. This suggests that CCTα not only displays structural but also functional adaptations to deep water conditions. The CCT complex is involved in the folding of ∼10% of newly synthesized proteins and has previously been categorized as ‘cold-shock’ protein in numerous eukaryotes. We thus propose that adaptation mechanisms to cold and deep-sea environments may be linked and highlight that efficient protein biogenesis, including protein folding and translation, are key metabolic deep-sea adaptations.

## Introduction

The deep ocean (>200m) covers approximately two-thirds of the global sea floor area, yet it is among the least studied environments of our planet in terms of biodiversity, habitats and ecosystem functioning (Ramirez-Llodra et al. 2010). It harbors specific environmental conditions such as high hydrostatic pressure (on average ∼400 atm), low temperatures (0-4°C), absence of light and scarcity of food. Life at great depths requires multiple metabolic adaptations resulting in a physiological bottleneck (Gross & Jaenicke 1994) limiting the vertical distribution of species (Brown & Thatje 2014; Somero 1992). It has also been proposed that high pressure may drive speciation events (Brown & Thatje 2014; Gaither et al. 2016). Numerous cellular processes such as enzymatic processes, protein folding, assembly of multi-subunit proteins and lipoprotein membranes are impacted by high pressure and low temperature (Carney 2005; Jaenicke 1991; Pradillon & Gaill 2007; Somero 1992). More specifically, cellular mechanisms of deep-sea adaptation have been recently reviewed (Yancey 2020) and include changes in membrane fluidity (high pressure and low temperature rigidify membranes), extrinsic adaptations (i.e. changes in the cellular milieu such as the expression of molecular chaperones or increased concentration of piezolytes, i.e. stabilizing solutes), and intrinsic adaptations (i.e. changes in protein sequences). An essential intrinsic adaptation of deep-sea proteins is their increased stability (i.e. more resistant to denaturation) compared to their shallow-water counterparts (Gross & Jaenicke 1994; Siebenaller 2010; Somero 1992, 1990, 2003). This has been shown in numerous studies, which mainly focused on a handful of proteins (such as lactate dehydrogenase) in deep-sea fishes (Dahlhoff & Somero 1991; Lemaire et al. 2018; Morita 2008, 2003; Siebenaller & Somero 1979, 1978; Suka et al. 2019; Wakai et al. 2014). Interestingly, patterns of protein adaptation to temperature show higher flexibility (i.e. decreased stability) with decreasing temperature (Fields et al. 2015; Saarman et al. 2017). As pressure and temperature strongly co-vary in the deep sea - temperature decreases as pressure increases - it therefore can be difficult to disentangle the respective combined or opposing effects of these factors on protein stability evolution.

Patterns of positive selection have been investigated to uncover genes underlying adaptation to specific environments, including the deep-sea, in non-model species (Kober & Pogson 2017; Lan et al. 2018; Oliver et al. 2010; Sun et al. 2017; Zhang et al. 2017; Weber et al. 2017). Although valuable, these studies typically focus on a single or few shallow-deep transitions in a limited number of species, and thus lack the comparative power to separate confounding effects. With almost 2100 species, brittle stars (Ophiuroidea) are a large and ancient class of echinoderms (Stöhr et al. 2017, 2012). These diverse marine invertebrates have colonized every marine habitat, highlighting their strong adaptive abilities. Furthermore, their phylogeny is well-resolved (O’Hara et al. 2017, 2014) and they represent a major component of the deep-sea fauna in terms of abundance and species diversity (Christodoulou et al. 2019a; Christodoulou et al. 2019b), making them important models for marine biodiversity and biogeography (O’Hara et al. 2019; Woolley et al. 2016). It is usually assumed that deep-sea organisms colonized the deep-sea from shallow waters (Brown & Thatje 2014); however, colonization from deep to shallow waters has also been reported (Bribiesca-Contreras et al. 2017; Brown & Thatje 2014). Four large independent ophiuroid families (Amphiuridae, Ophiodermatidae, Ophiomyxidae and Ophiotrichidae) have a common ancestor from shallow water with extant species occurring in deep and shallow seas (Bribiesca-Contreras et al. 2017). Due to these repeated and independent deep-sea colonization events, these four brittle star families provide an ideal framework to test for convergent molecular evolution to the deep sea.

## Materials and Methods

### Phylogenomic data generation and processing

The data used in this study is an extension (436 additional samples) of a previously published exon-capture phylogenomic datamatrix of 1484 exons in 416 genes for 708 individual ophiuroid samples representative of the whole class Ophiuroidea (O’Hara et al. 2019). The full dataset used here encompassed 1144 individual ophiuroid samples accounting for 826 species. Details on specimen collection, environmental parameters and list of species are available in Table S1. The set of 416 single-copy genes were first determined in a phylogenomic analysis to resolve the phylogeny of Ophiuroidea (O’Hara et al. 2014). These genes were rigorously screened to be one-to-one orthologs in 52 ophiuroid transcriptomes (representative of the entire class), in the sea urchin *Stronlgylocentrotus purpuratus*, the hemichordate *Saccoglossus kowalevskii* and the fish *Danio rerio*. The subsequent exon-capture system laboratory, bioinformatic and phylogenetic procedures are described in previous works (O’Hara et al. 2017; Hugall et al. 2016) and summarized in dryad packages https://doi.org/10.5061/dryad.db339/10 and https://datadryad.org/stash/dataset/doi:10.5061/dryad.rb334. Briefly, base-calling used a minimum read coverage of five. Exon boundaries were initially based on the *Strongylocentrotus purpuratus* and *Danio rerio* genomes, and then revised using the exon-capture read mapping information. For all selection analyses, each codon immediately adjacent to exon boundaries was ignored. The primary data had IUPAC-coded heterozygous sites, which were then randomly resolved. However, these sites had little influence as both ambiguity-coded and randomly resolved datasets returned the same positive selection test results. A global phylogenetic tree of all 1144 samples for 416 genes (273kb sites) was generated via RAxML v.8.2.11 (Stamatakis 2014) using a codon position partition model. First a fully-resolved all-compatible consensus topology was generated from 200 RAxML fast bootstrap samples (the -f –d command), onto which branch lengths were then optimized using a codon position GTR-Γ model (the -f –e command). The tree was then rooted according to (O’Hara et al. 2017) defining the sister superorders Ophintegrida and Euryophiurida.

Four brittle star families were investigated that included species displaying independent events of deep-sea colonization from shallow-water (Bribiesca-Contreras et al. 2017): Amphiuridae (111 individuals from 95 species, depth range: -0.5m to -5193m; temperature range: -1.6°C to 28.8°C), Ophiodermatidae (60 individuals from 38 species, depth range: -0.5m to -1668m; temperature range: 2.6°C to 28.3°C), Ophiomyxidae (41 individuals from 29 species, depth range: -1.5m to -792m; temperature range: 4.6°C to 28.7°C) and Ophiotrichidae (78 individuals from 62 species, depth range: -1m to -405m; temperature range: 10.5°C to 29.5°C) (Figure 1A). Positive selection analyses were conducted separately per family. 1664 alignments were generated, representing each gene (416) in each family (4). In each alignment, a maximum of 30% missing data per sequence was allowed. Alignments that lacked deep species (>200m) after filtering were not used. As all these four families belong to the superorder Ophintegrida, sequences of *Asteronyx loveni* belonging to the sister superorder Euryophiurida (Asteronychidae) were used as outgroups. After filtering, 1649 alignments were available for further analyses.

**Fig. 1:**
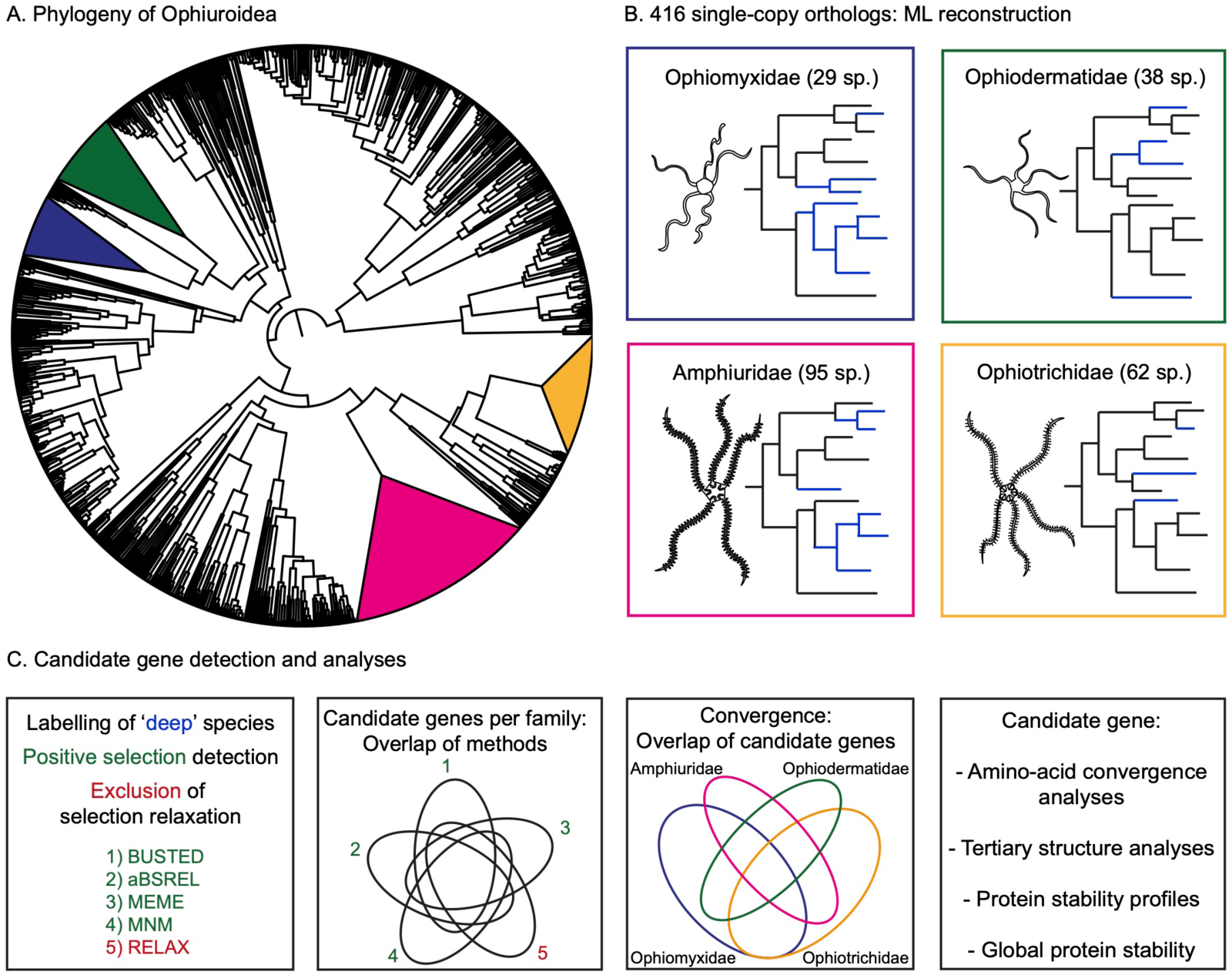
Workflow used in this study. A: Schematic representation of the phylogeny of Ophiuroidea (redrawn from Christodoulou et al. (2019b)). Four families (288 individuals from 216 species) with a shallow-water common ancestor and extant species in shallow (0-200m) and deep (>200m) environments are highlighted in different colors. The width of each triangle is proportional to the sampled number of species in each family. Ophiomyxidae (blue), Ophiodermatidae (green), Amphiuridae (pink) and Ophiotrichidae (yellow). B: For each family (number of species used in brackets) and each one of the 416 single-copy orthologs, Maximum Likelihood (ML) reconstructions were performed. Trees are drawn for illustration only with the branches of shallow-water (0-200m) species colored in black and deep-water (>200m) species in blue. C: For each resulting ML tree, deep species were labeled as foreground branches and four positive selection detection methods were used (BUSTED, aBSREL, MEME, MNM). To detect and exclude candidate genes displaying relaxation of selection, i.e. accumulation of substitutions not due to increased selection pressure, the method RELAX was used. The final set of candidate genes for each family encompassed genes positively selected in at least 3 methods and not displaying relaxation of selection. Convergent evolution was examined by overlapping candidate genes per family. For the most interesting candidate gene, amino-acid convergence and tertiary structure analyses were performed, and protein stability profiles and global protein stability were calculated.

### Phylogenetic reconstruction and positive selection analyses

For each of the 1649 alignments, a Maximum-Likelihood phylogeny was reconstructed using RAxML v.8.2.11 with the following parameters: -x 12345 -# 100 -f a -m GTRGAMMA -p 12345 (Figure 1B). Deep (>200m) species (tips) and monophyletic groups of deep species (nodes) were labeled as “Foreground” branches for positive selection analyses (Amphiuridae: 46 species; 11 independent events of deep-sea colonization; Ophiodermatidae: 7 species; 4 independent events; Ophiomyxidae: 10 species; 4 independent events; Ophiotrichidae: 4 species; 4 independent events). Then, the package HyPhy was used to conduct several positive selection analyses (Figure 1C): 1) BUSTED (Branch-site Unrestricted Statistical Test for Episodic Diversification) (Murrell et al. 2015) to test for gene-wide positive selection (at least one site on at least one branch); 2) aBSREL (adaptive Branch-Site Random Effects Likelihood) (Smith et al. 2015) to detect specific branches evolving under episodic positive selection; 3) MEME (Mixed Effects Model of Evolution) (Murrell et al. 2012) to find sites evolving under episodic positive selection. 4) MNM method; it has been shown recently that mutations at adjacent sites often occur as a result of the same mutational event (i.e. multinucleotide mutations, MNMs) and therefore may bias classical branch-site tests for positive selection (Venkat et al. 2018). The authors of that study developed a new model of positive selection detection incorporating MNMs, which we also used on deep lineages (>200m). For each gene, p-values were corrected for multiple testing using the Holm method (Holm 1979). The p-value significance level used for all the positive selection detection methods was 0.05. Finally, we used: 5) RELAX (Wertheim et al. 2014) to test for relaxation of selection, and exclude potential candidate genes displaying relaxation of selection. For each of the four families, positively selected candidate genes of each method were overlapped on a Venn diagram (Figures 1C and S1). To be considered as a candidate gene for positive selection in one family and to minimize the risk of false positives, a gene had to display a significant signal in at least three out of four methods including MEME and MNM (BUSTED, MEME, MNM or MEME, MNM, aBSREL) and not display relaxation of selection (RELAX). This set of candidates was used for functional annotation. Final sets of positively selected genes per family were then compared among each other to test for convergent evolution. To confirm that positive selection was detected only in deep-sea lineages, positive selection was also tested in shallow-water lineages for the five genes displaying convergent positive selection signatures (CCTα, PFD3, tkt, rpl34, rpl8; see Results, Tables 1, S3). For each family, the same number of shallow-water species as was used for deep-water species was randomly labeled as ‘Foreground’ in each gene tree and positive selection tests were performed for the five methods as described above.

**Table 1:**
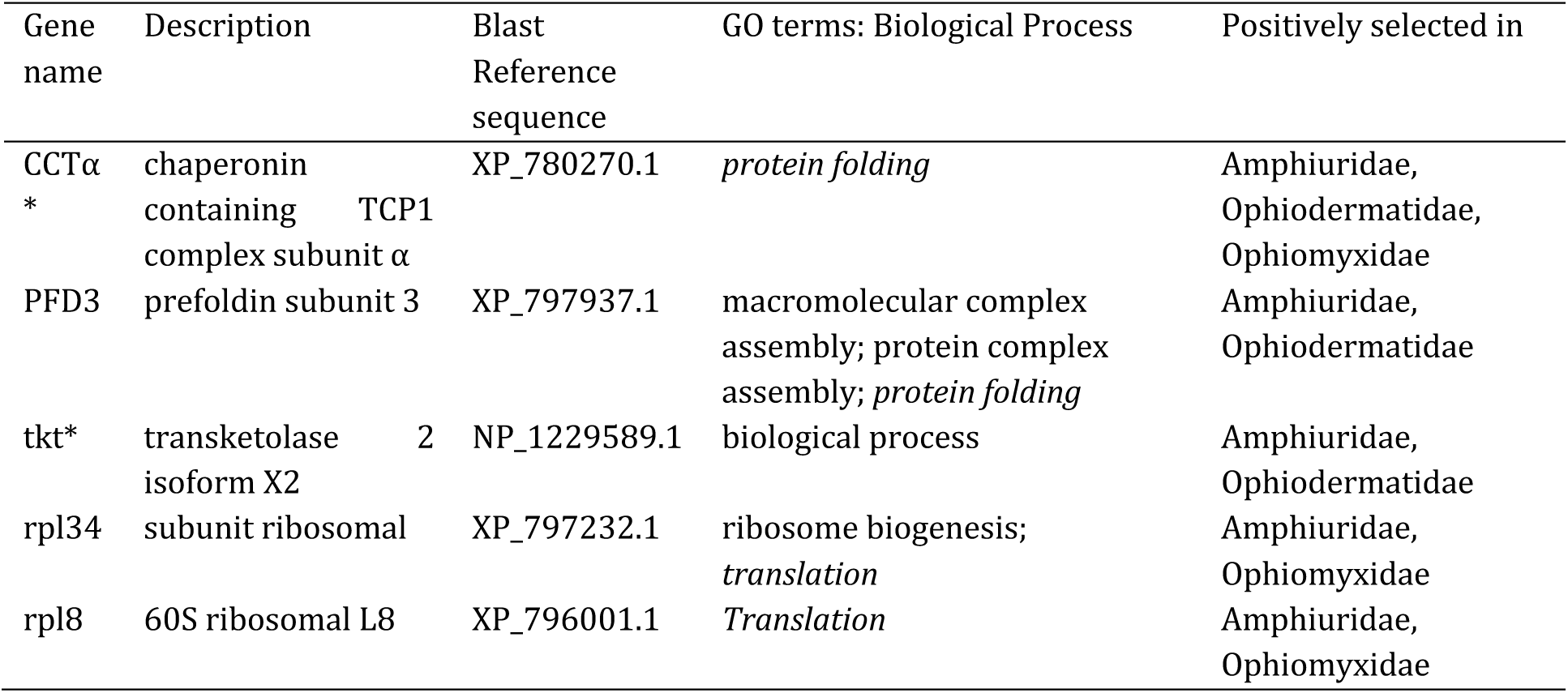
common positively selected candidate genes in three families and their characteristics (3 of 4 methods, not displaying relaxation of selection). *Positively selected in 4 of 4 methods, not displaying relaxation of selection. italic: common Biological Process annotation.

### Gene Ontology annotations and amino-acid convergence analyses

To explore which functions may be involved in deep-sea adaptation, the representative sequence of each of the 416 genes was extracted from the sea urchin *Strongylocentrotus purpuratus* genome and blasted against the nr database from NCBI using BLAST+. We used *S. purpuratus* as reference because sequence annotation for this species is of high quality (no high-quality brittle star reference genome is currently available) and to use a single complete representative sequence for each gene. The top 50 hits were extracted and loaded in BLAST2GO v.4.1. for annotation (Conesa et al. 2005). Mapping, annotation and slim ontology (i.e. GO subsets of broader categories) were performed with BLAST2GO using default parameters, except for the annotation cut-off parameter, which was set to 45. GO categories were described using the level 3 of slim ontology.

CCTα, the only candidate gene displaying positive selection signal in three families (see Results) was further analyzed for signatures of convergent evolution. Specifically, amino-acid profiles were investigated for convergent shifts using PCOC (Rey et al. 2018). This method, which has been shown to display high sensitivity and specificity, detects convergent shifts in amino-acid preferences rather than convergent substitutions. The CCTα amino-acid alignment encompassing the four families and outgroups was used to generate a maximum-likelihood phylogeny as previously described but this time using the PROTGAMMAWAG protein model of sequence evolution (Figure S2). For each family, positively selected branches resulting from aBSREL analyses were labeled as foreground branches (i.e. the branches with the convergent phenotype in the nucleotide topology) in four different scenarios: i) Amphiuridae, Ophiodermatidae, Ophiomyxidae; ii) Amphiuridae, Ophiodermatidae; iii) Amphiuridae, Ophiomyxidae; iv) Ophiodermatidae, Ophiomyxidae. Detection of amino-acid convergence in these four scenarios was then performed using PCOC and a detection threshold of 0.9 (Rey et al. 2018).

### Protein structure modeling, protein stability profile and overall protein stability

To infer the position of positively selected mutations on CCTα, the corresponding amino-acid sequence of the individual *Amphiura constricta* MVF214041 was used to obtain the secondary and tertiary protein structures of this gene. This shallow-water species was chosen because its CCTα sequence had no missing data. The secondary structure was modeled using InterPro 72.0 web browser (https://www.ebi.ac.uk/interpro/). The protein model was generated using the normal mode of the online Phyre^2^ server (http://www.sbg.bio.ic.ac.uk/~phyre2/) (Kelley et al. 2015). The online server EzMol 1.22 was used for image visualization and production (http://www.sbg.bio.ic.ac.uk/ezmol/) (Reynolds et al. 2018). The online server NetSurfP-2.0 (http://www.cbs.dtu.dk/services/NetSurfP/) was used to predict amino-acid surface accessibility and secondary structures.

We then examined the protein stability profiles of CCTα across the whole ophiuroid class (967 sequences with less than 30% missing sites, representing 725 species) using eScape v2.1 (Gu & Hilser 2009, 2008). This algorithm calculates a per-site estimate of Gibbs free energy of stabilization based on a sliding window of 20 residues. More specifically, it models the contribution of each residue to the stability constant, a metric that represents the equilibrium of the natively folded and the multiple unfolded states of a protein (D’Aquino et al. 1996). Sites adapted to elevated pressure (or high temperature at atmospheric pressure) are expected to display stabilizing mutations (i.e. more negative delta G values), whereas sites adapted to low temperatures at atmospheric pressure are expected to display mutations increasing flexibility (i.e. decreasing stability, thus more positive delta G values) (Fields et al. 2015; Saarman et al. 2017). For each site of the apical domain (codons 203-353), we calculated the average delta G value for all 324 shallow-water species (424 individuals) (0-200m) and 401 deep-water species (543 individuals) (>200m) and their respective standard deviation. To test the difference between these average values in a phylogenetic context, we used phylogenetically-corrected ANOVA (R function phylANOVA of the phytools v.0.6-60 R package; 10,000 simulations). To correct for relatedness among species, we used the global RAxML phylogenetic tree pruned to the 967 tips. To investigate regions rather than individual codons, we contrasted shallow vs. deep species along the whole gene, averaging delta G values across 10 residues and performing a phylogenetically-corrected ANOVA as previously described.

Finally, we tested how the mutations at the positively selected sites, and the ones with local stability values significantly different between shallow and deep species in the phylogenetically-corrected ANOVA, impacted the global protein stability. For each one of these sites, we identified ‘shallow-water’ and ‘deep-water’ mutations, and used the shallow-water Amphiuridae *Amphiura constricta* MVF214041 sequence as template to add these ‘shallow-water’ and ‘deep-water’ mutations one at the time. A model structure for each one of these new sequences was generated using the normal mode of the online Phyre^2^ server (http://www.sbg.bio.ic.ac.uk/~phyre2/) (Kelley et al. 2015). The global stability of each model structure was then calculated using the ‘stability’ function of the program FoldX with default values (Schymkowitz et al. 2005). For each site, the ΔΔG value between ‘deep-water’ (mutant) and ‘shallow-water’ (wild-type) mutations was calculated, where ΔΔG > 0.5 kcal/mol indicates stabilizing mutations, ΔΔG < -0.5 kcal/mol indicates destabilizing mutation, and -0.5 kcal/mol < ΔΔG < 0.5 kcal/mol indicates neutral mutations (Khatun et al. 2004; Khan & Vihinen 2010).

## Results

### Five genes involved in protein biogenesis are recurrently positively selected in deep-sea brittle stars

We used 416 single-copy orthologs from 216 species (288 individuals) of four brittle star families (Figure 1A) to examine patterns of positive selection in deep-sea species (>200m). For each gene of each family, we used four different positive selection methods and one method detecting relaxation of selection (Figure 1B-C). Given that our dataset is not representative of the whole genome, we chose a stringent approach to focus on a few candidate genes displaying strong signals of positive selection. Therefore, we only kept candidate genes that had significant signature of positive selection in at least three methods and which also did not show relaxation of selection. We found 36 candidate genes in Amphiuridae, 9 in Ophiodermatidae, 6 in Ophiomyxidae and none in Ophiotrichidae (Table S2; Figure S1). Four genes involved in embryo development were positively selected in Amphiuridae. Common functions among candidate genes from different families include protein folding, translation (i.e. ribosomal proteins), catabolic process and cell proliferation (Table S2).

Five genes were positively selected in at least two families, which are notably involved in protein folding and in translation (Table 1). Among these, CCTα was positively selected in all three families and significant in each one of the selection detection methods (Table 1). To confirm that positive selection was detected only in shallow-deep transition, we performed positive selection analyses on these five common candidate genes but this time labelling shallow-water lineages as ‘Foreground’. Most of the genes did not display signatures of positive selection in shallow-water environments, except PFD3 in Amphiuridae and tkt in Ophiotrichidae, which were significant in the BUSTED, aBSREL and MEME methods (Table S3). This suggests that these two genes are evolving faster not only in response to deep-sea conditions but also possibly in response to additional environmental conditions.

CCTα, the sole candidate gene detected as significantly positively selected in three families in the shallow-deep contrast, did not display signatures of positive selection in shallow-water environments (Table S3). It is a subunit of the octameric Chaperonin Containing TCP1 (CCT) complex, a cytosolic eukaryotic chaperonin having a central role in protein folding (Figure 2A-D) (Bueno-Carrasco & Cuéllar 2019; Valpuesta et al. 2005). The three other CCT subunits present in our dataset, CCTδ, CCTε and CCTη, did not display consistent signatures of positive selection; rather, while positive selection was detected at the site level in all subunits, relaxation of selection was detected in all 3 subunits in Amphiuridae (Table S6), indicating that the accumulation of mutations in these subunits is not clearly due to natural selection. Therefore, CCTα was the only one out of four subunits to show consistent signatures of positive selection across methods. The remaining four subunits of the CCT complex could not be tested as they were not present in our dataset.

**Fig. 2:**
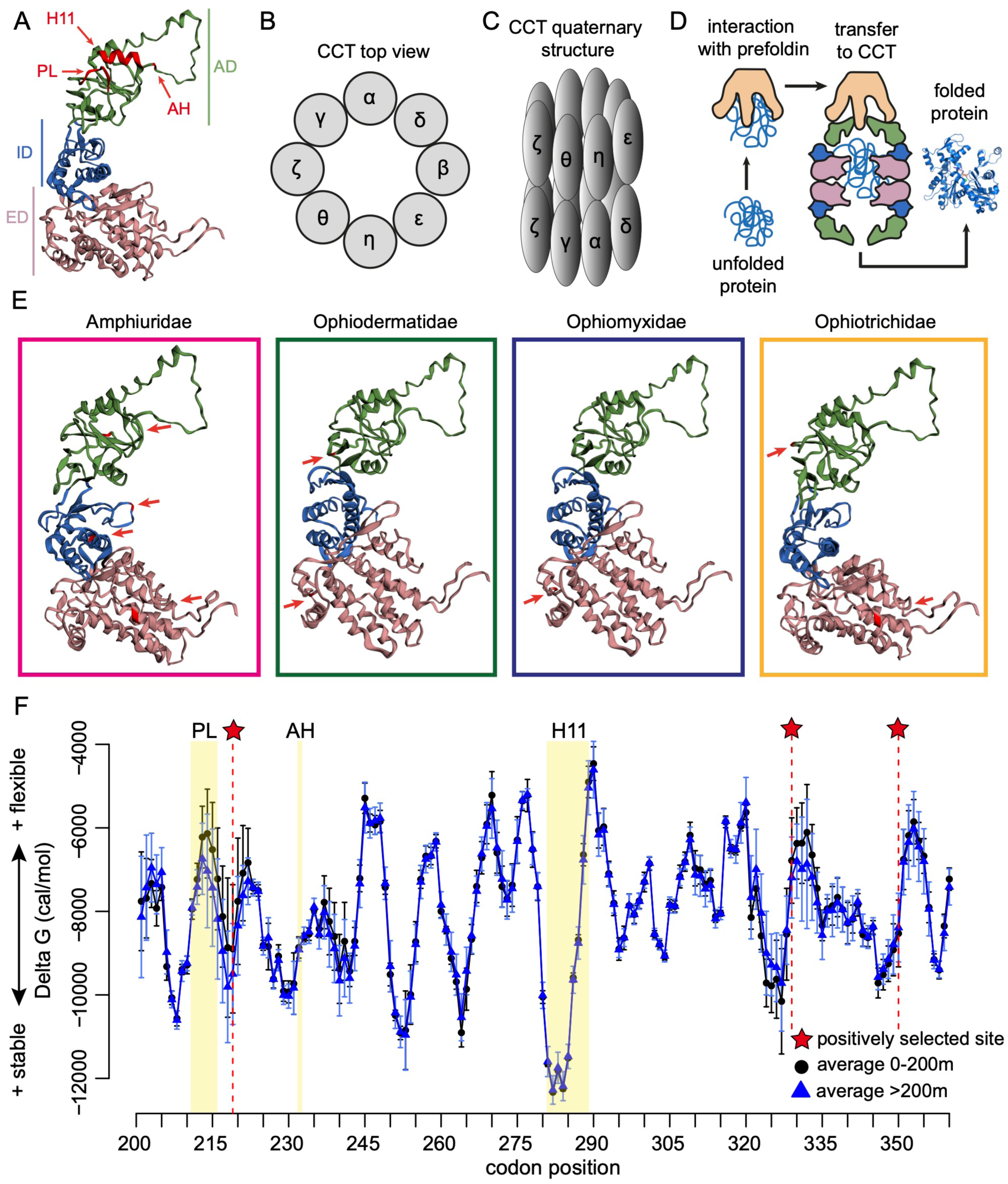
Structure and function of the CCT complex, selection analyses on CCTα and comparison of local stability values from CCTα apical domain between shallow and deep species. A: Model of tertiary structure of the CCTα subunit. Each subunit is composed of an apical domain (AD; green) containing the substrate binding regions (PL: Proximal Loop; AH: Apical Hinge; H11: Helix 11), an intermediate domain (ID; blue) and an equatorial domain (ED; pink) containing the ATP binding sites and where hydrolysis takes place. B: Model of the top view of the CCT complex, encompassing 8 paralogous subunits. C: Quaternary structure model of the CCT complex encompassing a double ring of 8 paralogous subunits. D: Simplified model of Prefoldin (PFD)-CCT interaction in the folding of newly synthesized actin or tubulin. A-D: Adapted from Bueno-Carrasco & Cuellar, 2018, “Mechanism and Function of the Eukaryotic Chaperonin CCT”. E: Localization of the positively selected sites on the tertiary structure of CCTα in the four ophiuroid families investigated. F: Average protein stability profiles and respective standard deviations (vertical bars) for each codon of the CCTα apical domain in 324 species (424 individuals) from shallow water (0-200m; in black) and 401 species (543 individuals) from deep water (>200m; in blue) representative of the whole ophiuroid class. A smaller (i.e. more negative) value of delta G is indicative of substitutions increasing stability. The substrate binding regions PL, AH and H11 are highlighted as well as the positively selected sites.

Interestingly, PFD3, a subunit of the hexameric co-chaperone prefoldin interacting with CCT (Gestaut et al. 2019; Martín-Benito et al. 2002) was positively selected in two families (Amphiuridae: MNM, MEME, BUSTED; Ophiodermatidae: MNM, aBSREL, MEME, BUSTED) (Table 1; Tables S2, S6; Figure 2D), although it was also positively selected in shallow-water Amphiuridae (Table S3). The two other prefoldin subunits present in our dataset (PFD1 and PFD5) did not show a consistent signature of positive selection (Table S4), suggesting that sub-units of this co-chaperone can evolve relatively independently from each other. Finally, two ribosomal proteins (Rpl8 and Rpl34) were recurrently positively selected in Amphiuridae and Ophiomyxidae, while the ribosomal protein S6 was positively selected in Ophiodermatidae (Table 1; Table S2).

### CCTα and deep-sea adaptation: convergence at the gene but not amino-acid level

While positive selection was detected at the gene level in three families, positive selection at the site level (MEME) was detected in all four families. The sites displaying positive selection in CCTα were not the same among the four families, except for site 26, which was shared between Amphiuridae and Ophiotrichidae (Figure 2E; Table 2; Table S5). Three sites were found in the equatorial domain, i.e. the ATP binding region, two sites were found in the intermediate domain and three sites were found in the apical domain, i.e. the substrate binding region (Figure 3) (Bueno-Carrasco & Cuéllar 2019). Furthermore, we found that 7 out 8 positively selected sites increased the global protein stability, suggesting that these sites may be involved in structural adaptation of CCTα (Table 2). This is corroborated by the fact that 6 out 8 mutations involve changes towards hydrophobic amino-acids, which are known to increase protein stability (Kellis et al. 1988; Pace et al. 2011). Finally, convergent evolution at the site level was not detected when examining amino-acid profiles (PCOC posterior probabilities at all sites < 0.9).

**Table 2:** Summary of the positively selected sites in each family and sites with local stability values significantly different between shallow and deep species. For each site, the ‘deep’ and ‘shallow’ amino-acids are reported, as well as their impact on global protein stability: ΔΔG>0.5 kcal/mol: mutations increasing stability; ΔΔG<-0.5 kcal/mol: mutations decreasing stability; -0.5 kcal/mol <ΔΔG<0.5 kcal/mol: neutral mutations in terms of stability. ED: equatorial domain; ID: intermediate domain; AD: apical domain; PS: positively selected. *: the most common amino-acids between shallow and deep species are reported.

| Site | Tertiary structure | Detection method | Amino-acid ‘deep’ | Amino-acid property | Amino-acid ‘shallow’ | Amino-acid property | $\Delta\Delta G$ [kcal/mol] |
| --- | --- | --- | --- | --- | --- | --- | --- |
| 26 | ED | PS Amphiuroidae & PS Ophiotrichidae | A | hydrophobic | S | Polar uncharged | 34.7 |
| 175 | ID | PS Amphiuroidae | Y | hydrophobic | S | Polar uncharged | 11.8 |
| 203* | AD | phylANOVA | P | special case | Q | Polar uncharged | -17.3 |
| 215* | AD | phylANOVA | V | hydrophobic | P | special case | 2.1 |
| 219 | AD | PS Ophiotrichidae | I | hydrophobic | T | Polar uncharged | 3.7 |
| 263* | AD | phylANOVA | H | Positively charged | Q | Polar uncharged | 22.9 |
| 327* | AD | phylANOVA | S, V | Polar uncharged, hydrophobic | A | hydrophobic | 11.9, 30.1 |
| 328* | AD | phylANOVA | T | Polar uncharged | S | Polar uncharged | 27.6 |
| 329 | AD | PS Amphiuroidae | V | hydrophobic | M | hydrophobic | 15.7 |
| 350 | AD | PS Ophiidermatidae | I | hydrophobic | K | Positively charged | 15.7 |
| 375 | ID | PS Amphiuroidae | V | hydrophobic | I | hydrophobic | 12.3 |
| 462 | ED | PS Ophiomyxidae | C | special case | A | hydrophobic | -3.8 |
| 465 | ED | PS Ophiidermatidae | F | hydrophobic | K | Positively charged | 22.9 |

**Fig. 3:**
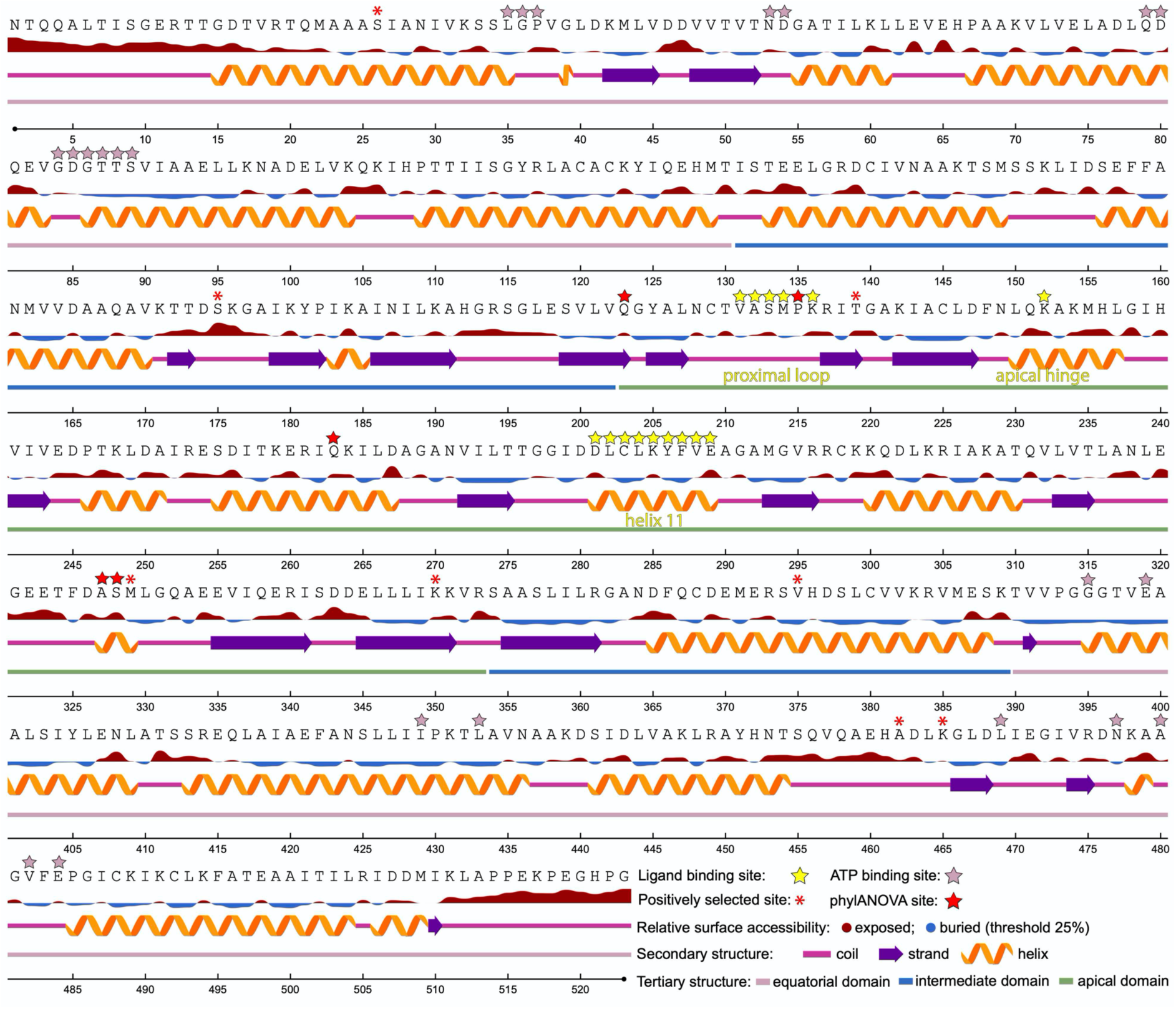
Primary structure of CCTα. Ligand binding sites (proximal loop, apical hinge and helix 11), ATP binding sites, positively selected sites (method MEME) and sites displaying different local stability values between shallow and deep species (method phylANOVA) are highlighted. The sequence of the shallow-water Amphiuridae *Amphiura constricta* is used as template.

### Energetic landscapes reveal structural adaptation within and next to the substrate binding region

Next we calculated site-specific protein stability profiles of CCTα in 967 individuals of 725 species representative of the whole Ophiuroidea class, to test the hypothesis that deep-sea adapted proteins are more stable than their shallow-water counterparts. For each site, we compared the average stability measure of 324 shallow-water species (0-200m depth) vs. 401 deep-water species (>200m depth), where lower ΔG values correspond to higher stability (Figures 2F, S3A). We focused on the apical domain as it encompasses the substrate binding region, whose position and structure are highly conserved across eukaryotes (Joachimiak et al. 2014). This region is composed of the proximal loop (PL), the apical hinge (AH) and Helix 11 (H11) (Figures 2A, 3). While AH and H11 are almost invariant across all ophiuroids, the stability measure was lower (i.e. more stable) in deep compared to shallow species within the PL and in two sites following the PL (codons 214-217), close to a positively selected site (codon 219) (Figures 2F, S3A). In contrast, three codons displayed significantly higher flexibility in deep compared to shallow species (Figure S3A,B), suggesting that increased flexibility may play a role in deep-sea adaptation outside the ligand binding region, possibly related to low temperature. Nevertheless, when averaging ΔG values across 10 codons, only the signal close to PL remained significant in the phylogenetically-corrected ANOVA contrasting stability values of shallow and deep species (Figure S4). This indicates that substitutions towards a more stable PL occurred independently in the ophiuroid tree of life. Finally, we examined how these mutations impacted the global (rather than local) stability value of the protein. We found that 4 out of 5 mutations increased the global protein stability, including the site located in the substrate binding region (i.e. site 215 in the PL), confirming the results found with the local stability values.

## Discussion

### Protein biogenesis is essential for deep-sea adaptation

In this study, we examined patterns of positive selection in 416 genes from four brittle star families representing replicated instances of deep-sea colonization. We found that four genes involved in functions related to protein biogenesis including protein folding and protein synthesis (i.e. ribosomal proteins) were convergently positively selected in three out of four families. Furthermore, an additional ribosomal protein was positively selected in Ophiodermatidae and two transcription factors were positively selected in Amphiuridae. Previous studies have shown that several genes involved in genetic information processing such as protein folding, transcription or translation (including ribosomal proteins) were positively selected in the hadal (>6000m depth) amphipod *Hirondella gigas* (Lan et al. 2017), in the bathyal (200-3000m depth) fish *Aldrovandia affinis* (Lan et al. 2018) and in the bathyal barnacle *Glyptelasma gigas* (Gan et al. 2020). Taken together, these and previous results indicate that adaptations in protein biogenesis are essential to deep-sea life.

At the individual family level, four genes related to embryo development were positively selected in Amphiuridae. Interestingly, nine genes related to the same function (embryo development; GO: 0009790) were detected as candidate genes for deep-sea adaptation between populations of the deep-sea fish *Coryphaenoides rupestris* living at different depths (Gaither et al. 2018). This suggests that there might be similar selection pressures on development at both young (i.e. species level) and old (i.e. family level) evolutionary timescales for life at great depths. Finally, no candidate genes for deep-sea adaptation were found in Ophiotrichidae across different positive selection methods. This might be explained by the fact that this family is the youngest from a fossil and a phylogenetic point of view (O’Hara et al. 2017). In that sense, it started colonizing the deep-sea later than the other families, and therefore there was possibly less time for natural selection to act. In support with this, the maximum sampling depth for Ophiotrichids was 405m, whereas it was deeper for the three other families (Amphiuridae: 5193m; Ophiodermatidae: 1668m; Ophiomyxidae: 792m).

### Independent evolutionary histories of subunits from macromolecular complexes

CCTα and prefoldin 3 (PFD3), the two genes recurrently positively selected involved in protein folding, are subunits of octameric and hexameric complexes, respectively. The CCT complex is estimated to fold ∼10% of newly synthesized proteins, including actin and tubulin, and is involved in numerous core cellular processes such as cytoskeleton formation, cell signaling, cell recognition and protein degradation (Bueno-Carrasco & Cuéllar 2019), while the prefoldin complex is known to assist CCT in actin folding (Martín-Benito et al. 2002). We tested four subunits of the octameric CCT complex, yet CCTα was the only one to show consistent signatures of positive selection across different methods. Indeed, the other tested subunits displayed relaxation of selection or no signatures of positive selection. This might be due to the different degrees of subunit specialization in CCT, as CCTα has intermediate binding properties (i.e. neither high ATP affinity nor high substrate affinity) compared to the other subunits (Bueno-Carrasco & Cuéllar 2019). Therefore, it is possible that CCTα can evolve more rapidly because it may experience less negative selection than the other subunits. In the same way, we tested three subunits of the hexameric prefoldin complex, and PFD3 was the only subunit to be consistently positively selected. While we acknowledge that we analyzed an existing dataset and thus did not test all subunits of both macromolecular complexes (4/8 in CCT; 3/6 in prefoldin), it appears surprising that individual subunits of macromolecular complexes can evolve relatively independently from each other.

We note that a previous study actually highlighted evolutionary independence of CCT subunits across old evolutionary timescales (i.e. among eukaryotes), showing that CCTα, CCTγ and CCTζ evolved under positive selection after duplication events which led to sub-functionalization in eukaryotes, most likely in response to folding increasingly complex cytosolic proteins (Fares & Wolfe 2003). Furthermore, it has been shown that the exact same mutation in the conserved ATP-binding region of all eight CCT subunits have dramatically different phenotypic outcomes in yeast, such as differential temperature sensitivity, differential growth in presence of actin and tubulin polymerization inhibitors, or differential over and under-expression of genes (Amit et al. 2010). This not only indicates that the functional activity of each subunit is not redundant but also possibly highlights ‘moonlighting’ proteins (i.e. other functions of individual subunits that are carried out not as part of the full complex) (Jeffery 2009; Huberts & van der Klei 2010). In that sense, the functional coherence of several hetero-oligeromeric protein complexes including CCT has been reviewed (Matalon et al. 2014). Interestingly, they found a lack of coherence among subunits at virtually all functional levels, including different levels of subunit abundance, differences in transcriptional regulation, differential subunit duplication or loss, and finally lack of coherence in genetic interaction. Explanations for this lack of coherence include enhancing complex assembly or subunit multi-functionality (Matalon et al. 2014). There is increasing evidence for the latter case, as many CCT monomer functions have been reported and recently reviewed (Vallin & Grantham 2019). Taken together, our results are consistent with the hypothesis that CCTα and PFD3 may be moonlighting proteins involved in deep-sea adaptation.

### Convergent evolution, functional and structural adaptations in CCTα

Our results suggest that convergent evolution was detected at the functional (i.e. protein folding) and at the gene levels (i.e. CCTα, PFD3) in deep-sea adaptation. Yet, it was not detected at the amino-acid level in CCTα, as the majority of positively selected sites were different among families, except for one site shared between Amphiuridae and Ophiotrichidae. This might be due to the old divergence time among the investigated brittle star families and the broad nature of the selection regime (i.e. depth and temperature). It has been shown that rates of molecular convergence decrease with time (Storz 2016) and the last common ancestor of Amphiuridae, Ophiodermatidae and Ophiomyxidae is estimated to be approximately 250 million years old (O’Hara et al. 2017). Furthermore, convergence at the amino-acid level is often less common than convergence at higher levels of biological hierarchy (e.g. gene, pathway or species levels) (Bolnick et al. 2018; Tenaillon et al. 2016, 2012).

Our results indicate that CCTα evolved under positive selection during deep-sea colonization and speciation events. These mutations possibly reveal functional adaptations of CCTα, such as adaptations in ATP or substrate binding. Indeed, it has previously been reported that the shallow groove created by the conserved Helix 11 and the flexible Proximal Loop allows the binding of a variety of substrates (Joachimiak et al. 2014; Yam et al. 2008), suggesting that the detected substitutions in the PL and in adjacent amino acids allow efficient substrate binding in deep-sea species. Similarly, in a study on metabolic enzymes from 37 ctenophores, numerous sites associated with adaptation to depth, temperature or both were located close to the substrate binding region (Winnikoff et al. 2019). A major substrate of the CCT complex and its co-chaperone prefoldin is actin, an essential cytoskeleton protein (Martín-Benito et al. 2002). While it should be noted that CCTα is not directly involved in actin folding (actin binds the subunits CCTδ, CCTβ and CCTε) (Llorca et al. 1999, 2000), examination of cytoskeletal proteins is of interest due to their essential role in cells. Unfortunately, as actin and tubulin were not included in our dataset, no conclusions about cytoskeletal proteins of deep-sea brittle stars can yet be drawn. However, actin of deep-sea fishes has been extensively studied, showing that it displays structural adaptations to high-pressure enhancing stability (Morita 2003; Koyama & Aizawa 2008; Wakai et al. 2014; Yancey 2020). Furthermore, genome analysis of the hadal fish *Pseudoliparis swirei* revealed that a gene family related to cytoskeleton was expanded, and several genes from three gene ontology categories linked to cytoskeleton were evolving under positive selection (Wang et al. 2019). Finally, 13 genes involved in cytoskeleton systems were positively selected in the deep-sea fish *A. affinis* (Lan et al. 2018), suggesting that cytoskeletal proteins are pivotal for deep-sea adaptation.

Furthermore, we tested the hypothesis that CCTα is structurally adapted to deep-sea conditions, as a common feature of high-pressure adapted proteins is to display increased stability (Somero 2003; Ohmae et al. 2013; Yancey & Siebenaller 2015). We investigated local and global stability in CCTα and found that four out of five sites with phylogenetically-corrected significantly different local stability showed increased global protein stability, suggesting that these mutations reflect structural deep-sea adaptation. Moreover, seven out of eight mutations at positively selected sites increased global protein stability, and involved hydrophobic amino-acid substitutions, which are known to increase protein stability (Kellis et al. 1988; Pace et al. 2011). It has also been shown that several proteins from the abyssal fish *Coryphaenoides armatus* displayed increased protein stability as an adaptation to hydrostatic pressure (Morita 2003, 2008; Brindley et al. 2008; Lemaire et al. 2018). Finally, several critical mutations in both the enzyme synthesizing trimethylamine oxide (TMAO: a heavily studied stabilizing cosolute or piezolyte (Yancey 2020)) and in HSP90, have been proposed to enhance overall protein stability in the hadal fish *Pseudoliparis swirei* (Wang et al. 2019).

### Role of molecular chaperones in deep-sea adaptation

We have shown that over deep evolutionary timescales, CCTα displays recurrent signatures of accelerated evolution and structural adaptation in transition from shallow to deep-sea habitats in brittle stars. This contrasts with previous findings on sea urchins where, CCTε -but not the other subunits-showed signatures of positive section, yet not in the two deep-sea species included in the work (Kober & Pogson 2017). A functional equivalent of the eukaryotic chaperonin CCT in bacteria is the chaperonin complex GroEL/GroES (Gupta 1990; Mayhew et al. 1996). This chaperonin has been shown to be constitutively highly expressed in the deep-sea bacteria *Shewanella*, and that a major target of this chaperonin was a subunit of the 50S ribosome (Sato et al. 2015). Furthermore, convergent evolution in chaperones has previously been shown (Draceni & Pechmann 2019), and there is evidence that other groups of molecular chaperones such as HSP70 and HSP90 (Kim et al. 2013) are involved in deep-sea adaptation (Yancey 2020). For instance, the HSP70 gene family was expanded in a deep-sea mussel (Sun et al. 2017) and convergent amino-acid changes have been reported in four out of five copies of the HSP90 protein in a hadal fish (Wang et al. 2019). In our dataset, two HSP90 genes were present (HSP90b1 and HSP90ab1), yet neither showed signatures of accelerated evolution. Nevertheless, only the examination of whole genomes will give a complete view of HSP family evolution in brittle stars, which should be explored in further studies.

### Similarities between deep-sea and cold adaptation mechanisms

There are numerous studies on the role of CCT at shorter evolutionary timescales, which reveal its role in cold-stress response. Notably, CCT has been characterized as a ‘cold-shock’ protein in several eukaryotes due to the overexpression of the investigated subunits when organisms were exposed to cold stress (He et al. 2017; Kayukawa et al. 2005; Somer et al. 2002; Yin et al. 2011). Furthermore, CCT has been shown to display specific structural (Pucciarelli et al. 2006) and functional (Cuellar et al. 2014) adaptations to cold environment in Antarctic fish, in addition to being also overexpressed in Antarctic fish exposed to heat stress (Buckley & Somero 2009). Moreover, there is evidence for a link between cold-stress response and high-pressure stress response in bacteria (Welch et al. 1993; Wemekamp-Kamphuis et al. 2002), although it has also been shown that heat-shock proteins are also expressed under high-pressure stress in bacteria (Aertsen et al. 2004) or constitutively in the deep-sea bacteria *Shewanella* (Sato et al. 2015). Additionally, transcriptome analyses revealed that cold-inducible protein families are expanded in a hadal amphipod (Lan et al. 2017), and several proteins involved in cold shock have been shown to evolve under positive selection in deep-sea amphipod and fish (Lan et al. 2018). Taken together, our findings support the hypothesis that cold shock proteins play an important role in deep-sea adaptation (Brown & Thatje 2014), which was also proposed by others (Lan et al. 2018).

### Limitations and outlook

We acknowledge that our study lacks functional validation to demonstrate that the changes are truly adaptive (which would be experimentally demanding as CCT folds ∼10% of newly synthesized proteins and we analyzed the sequences of more than 700 species), yet we minimized false inferences by applying stringent positive selection detection criteria. Furthermore, we used a *proxy* of functional validation by investigating *in silico* local and global protein stability in a dataset with great comparative power, both in terms of phylogenetic and environmental diversity. Finally, direct experimental testing on deep-sea organisms remains technically challenging, so we made use of the power of molecular data to indirectly reveal new insights in deep-sea adaptation. Further studies should include whole genomes to obtain a more complete view of deep-sea adaptation hotspots and possible mechanisms, such as recent studies on deep-sea fishes (Gaither et al. 2018; Wang et al. 2019), hydrothermal vent mussel (Sun et al. 2017) or cold seep gastropod (Liu et al. 2020). Also, while we focused on intrinsic adaptations (i.e. at the genomic level), mechanisms of extrinsic adaptations (i.e. changes in the cellular milieu) through, for instance, osmolyte concentration (piezolytes such as TMAO) or membrane phospholipid composition changes, should not be overlooked (Yancey & Siebenaller 2015; Somero 2003; Yancey 2020). However, this is beyond the scope of this study. With increasing interests in deep-sea biodiversity, ecosystems and resources in the last decades (Danovaro et al. 2017, 2014; Glover et al. 2018), these are exciting times for diving deeper into mechanisms of deep-sea adaptation.

## Acknowledgements

We are grateful to W. Salzburger for providing access to the HPC sciCORE cluster and to J. Sarrazin and S. Arnaud-Haond for comments on a previous version of this manuscript. We thank L. Bribiesca-Contreras for help with R analyses. We thank three anonymous reviewers and the editorial team for valuable comments on a previous version of this manuscript. Calculations were performed at sciCORE (http://scicore.unibas.ch/) scientific computing center at the University of Basel, Switzerland and at DATARMOR (http://www.ifremer.fr/pcdm) scientific computing center at the Pôle de Calcul et de Données Marines (PCDM), Ifremer, Brest, France. This work was supported by an Endeavour Postdoctoral Fellowship awarded by the Australian Department of Education and Training [grant number 6534_2018 to AATW] and a Marie Sklodowska-Curie Global Fellowship awarded by the European Union’s Horizon 2020 research and innovation programme [grant number 797326 to AATW].

## Data deposition

Phylogenomic data (including raw reads) and scripts for dataset generation are available in NCBI Bioproject PRJNA311384 and dryad packages: https://doi.org/10.5061/dryad.db339/10 and https://datadryad.org/stash/dataset/doi:10.5061/dryad.rb334.

## Supplementary figure captions

**Fig. S1:** Overlap of positively selected candidate genes among three brittle star families. A: Number of candidate genes per family positively selected in at least three positive selection detection methods and not displaying relaxation of selection. B: Number of candidate genes per family positively selected in all four positive selection detection methods and not displaying relaxation of selection. In both conditions of A and B, no gene was positively selected in the family Ophiotrichidae.

**Fig. S2:** Maximum-likelihood reconstruction of CCTα using amino-acid sequences. The four focal families (Amphiuridae, Ophiotrichae, Ophiomyxidae and Ophiodermatidae) and the outgroup (Asteronychidae) are labelled. Positively selected lineages (aBSREL method) of each family are highlighted in red.

**Fig. S3:** Comparison of local stability values from CCTα apical domain between shallow and deep species. A: Log transformed p-values of the phylogenetically-corrected ANOVA performed between average delta G values of shallow vs. deep species at each codon of the CCTα apical domain. The substrate binding regions PL, AH and H11 are highlighted in light orange. The positively selected sites are highlighted with a red star. P-value level corresponding to 0.05 is highlighted in red. The most significant codons in a “significance peak” (204, 214, 265 and 324) are highlighted. B: Beanplots of delta G values between shallow (0-200m) and deep (>200m) species for each one of the most significant codons in the phylogenetically-corrected ANOVA. Horizontal bar represents the average value of the dataset.

**Fig. S4:** Comparison of local stability values over 10-codon windows on the complete CCTα gene between shallow and deep species. A: Average protein stability profiles over 10 codon windows for the complete CCTα gene in 324 species (424 individuals) from shallow water (0-200m) and 401 species (543 individuals) from deep water (>200m) representative of the whole ophiuroid class. A smaller (i.e. more negative) value of delta G is indicative of substitutions increasing stability. The positively selected sites are highlighted with a red star. B: Log transformed p-values of the phylogenetically-corrected ANOVA performed between average delta G values over 10 codon windows of shallow vs. deep species. The positively selected sites are highlighted with a red star. P-value level corresponding to 0.05 is highlighted in red. C: Beanplots of delta G values between shallow (0-200m) and deep (>200m) species for both of the most significant 10 codon windows in the phylogenetically-corrected ANOVA. Horizontal bar represents the average value of the dataset.

## Supplementary tables (in separate excel file)

**Table S1:** List of species used in this study, GPS coordinates and environmental parameters at their sampling locations. Empty cells indicate missing data.

**Table S2:** Positively selected candidate genes per family and their Gene Ontology annotation (P: Biological Process; F: Molecular Function; C: Cellular Component). Genes positively selected in several families are highlighted in bold.

**Table S3:** Results of positive selection tests in shallow-water lineages for the five candidate genes for deep-sea adaptation. Significance level: 0.05. Significant results are in bold. NS: not significant

**Table S4:** Results of positive selection tests for the three prefoldin subunits for each family. Significance level: 0.05. Significant results are in bold. NS: not significant

**Table S5:** Sites displaying episodic positive selection in CCTα. Method used: MEME. Significance level: 0.05. Significant sites are in bold.

**Table S6:** Results of positive selection tests for the four CCT subunits for each family. Significant results are in bold. Significance level: 0.05. NS: not significant

