## Supplementary figures and images for "Convergent evolution and structural adaptation to the deep ocean in the protein folding chaperonin CCTα"

### Supplemental Figure 1

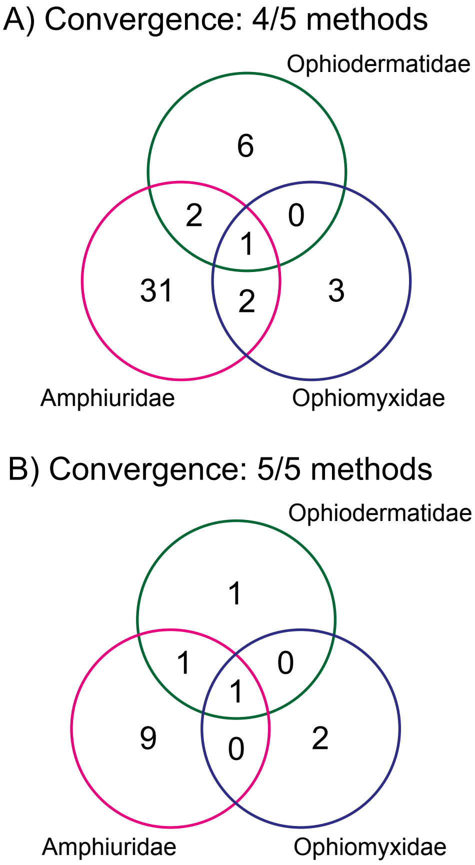

### Supplemental Figure 2

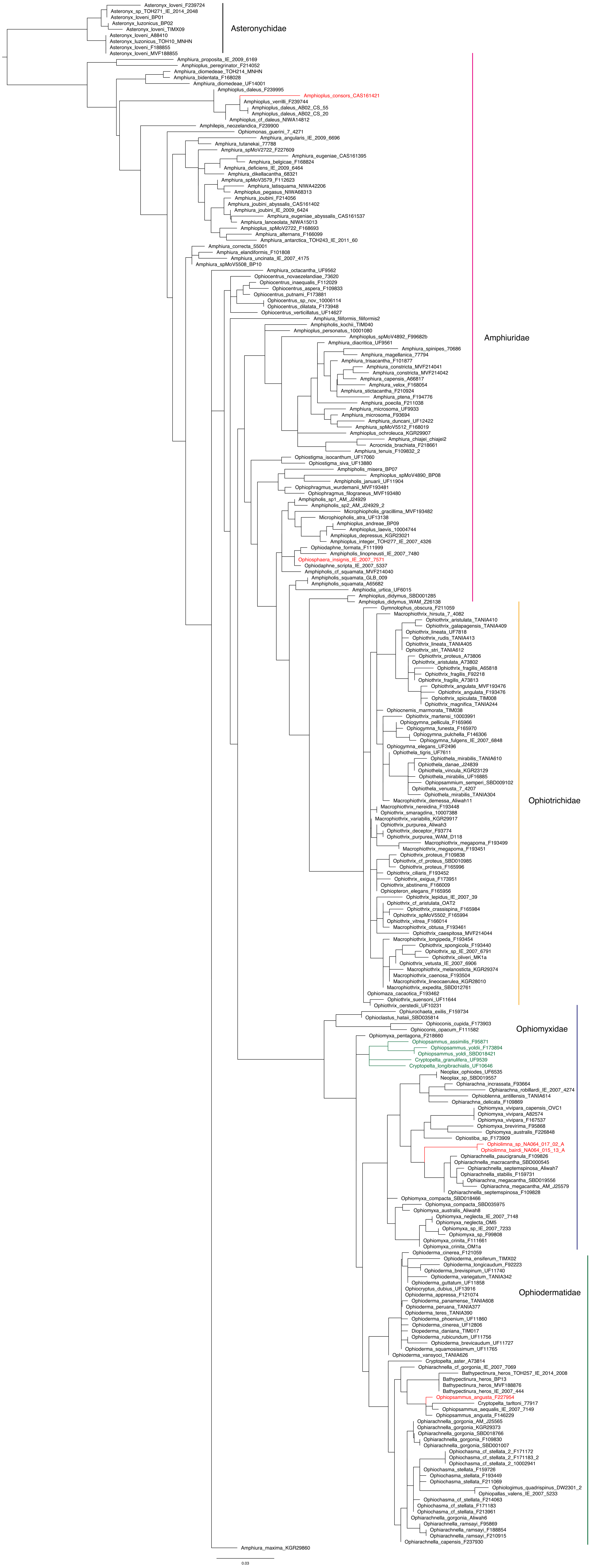

### Supplemental Figure 3

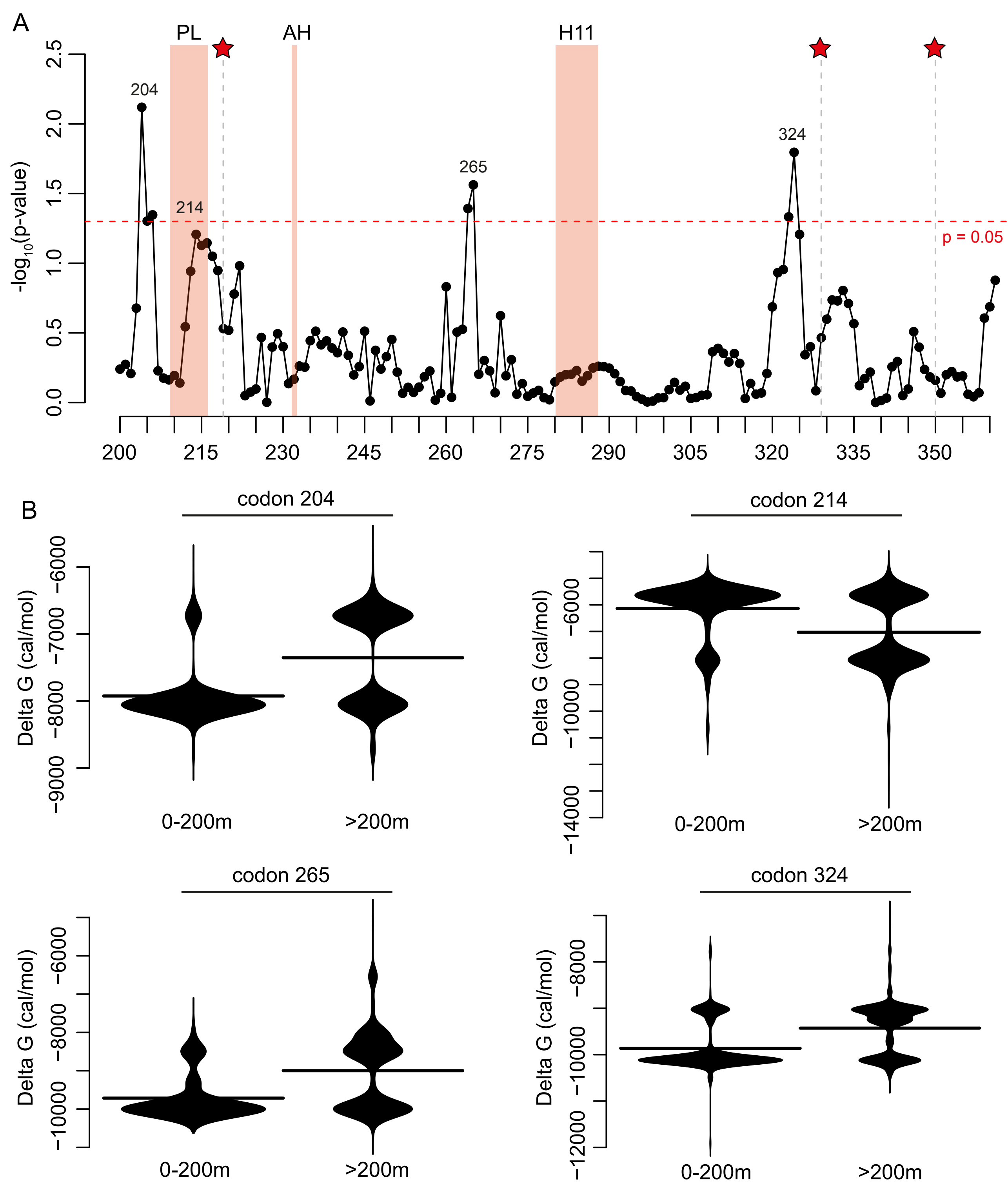

### Supplemental Figure 4

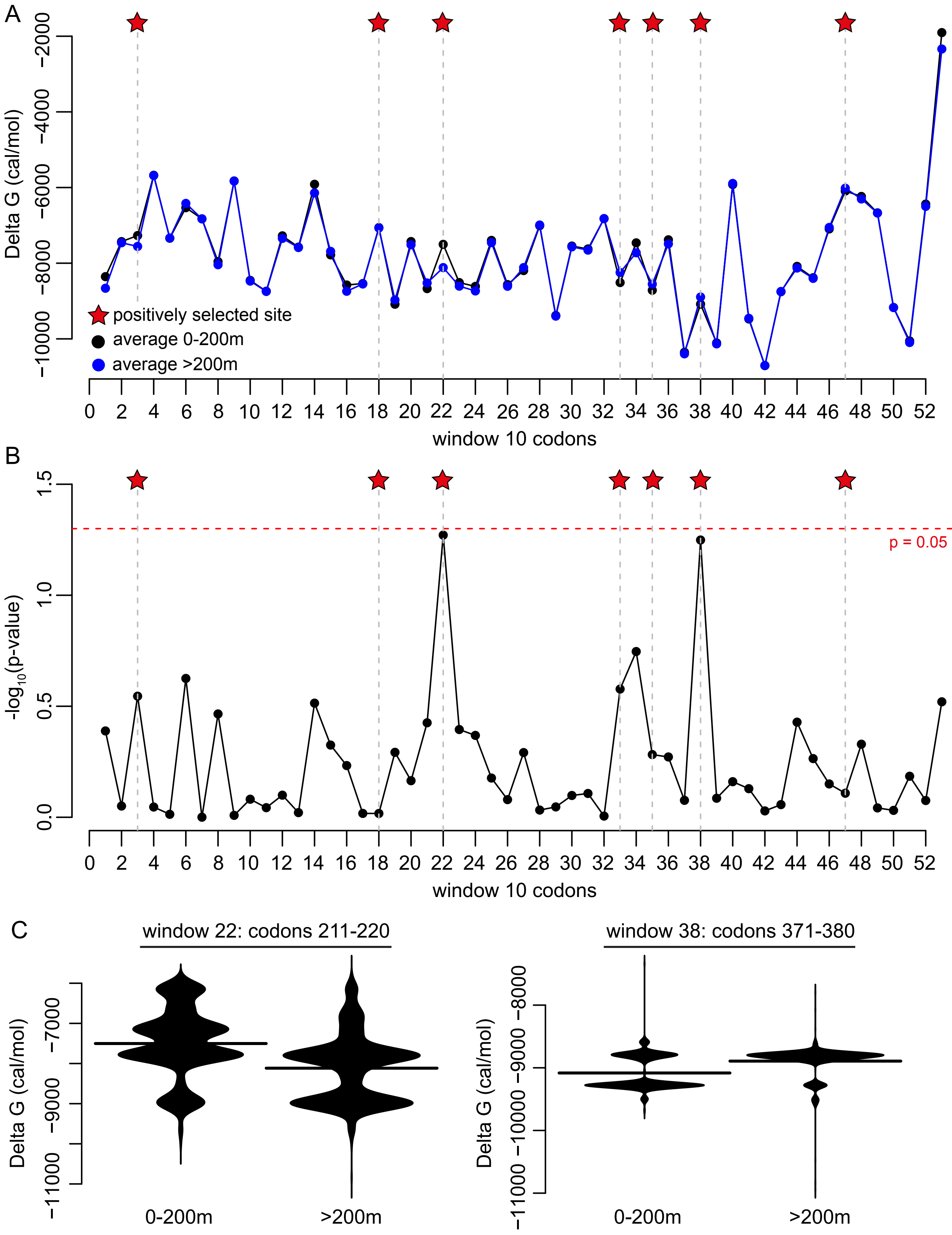
